# Revisiting post-stimulus theta activity: evidence for an aperiodic rather than oscillatory origin

**DOI:** 10.64898/2026.06.16.732609

**Authors:** Theo Vanneau, Michael Quiquempoix, Bradley Voytek, Mate Gyurkovics, Sophie Molholm

**Affiliations:** The Cognitive Neurophysiology Laboratory, Departments of Pediatrics & Neuroscience, Albert Einstein College of Medicine, Bronx, New York 10461, USA; Institut de Recherche Biomédicale des Armées (IRBA), 91223 Brétigny sur Orge, France; URP 7330 VIFASOM, Université Paris Cité, Hôtel Dieu, Paris, France; Department of Cognitive Science, UC San Diego, La Jolla, CA, USA; School of Psychology, University of East Anglia, Norwich, United Kingdom; The Frederick J. and Marion A. Schindler Cognitive Neurophysiology Laboratory, The Ernest J. Del Monte Institute for Neuroscience, Department of Neuroscience, University of Rochester School of Medicine and Dentistry, Rochester, New York 14642, USA

## Abstract

The aperiodic, 1/f-like component of electrophysiological activity is increasingly recognized as a meaningful feature of neural function, rather than background noise. In parallel, many EEG studies report transient changes in oscillatory power following stimulus onset and interpret these effects as signatures of attention, salience, or cognitive control. However, such conclusions usually rely on baseline normalization procedures that assume aperiodic activity remains stable from pre-to post-stimulus periods. Using high-density EEG recordings from typically developing children, we tested this assumption in two paradigms: an audiovisual simple reaction-time task (n = 36) and a visual oddball task (n = 38). For each task, conventional spectral analyses were compared with analyses that explicitly modeled and removed the aperiodic component in both pre- and post-stimulus windows. Across tasks, stimulus onset was associated with robust increases in aperiodic exponent and offset, indicating systematic changes in the 1/f component of the spectrum. In the audiovisual task, these changes were modality-specific, with central, parieto-occipital, or combined topographies depending on stimulus type. These effects were reduced but remained significant after ERP removal, indicating that they were not fully explained by phase-locked activity. Critically, once aperiodic activity was accounted for, the apparent post-stimulus increase in theta power was largely abolished in both tasks, including the canonical fronto-central theta enhancement to infrequent targets in the oddball paradigm. The conventional method also overestimated the magnitude of beta desynchronization, particularly in the induced (ERP-removed) signal. The apparent gamma desynchronization detected by conventional analyses was reversed after aperiodic correction, revealing either synchronization or no change, indicating that it reflects a spurious consequence of spectral slope steepening rather than a true suppression of gamma oscillatory activity. In contrast, alpha desynchronization remained robust after aperiodic correction and was in fact enhanced, suggesting it reflects genuine oscillatory suppression. Together, these findings indicate that a substantial portion of conventional time-frequency effects, particularly apparent theta synchronization, may reflect changes in aperiodic activity in response to stimulation rather than genuine periodic oscillations, challenging core assumptions of conventional time-frequency analyses.

Graphical abstract

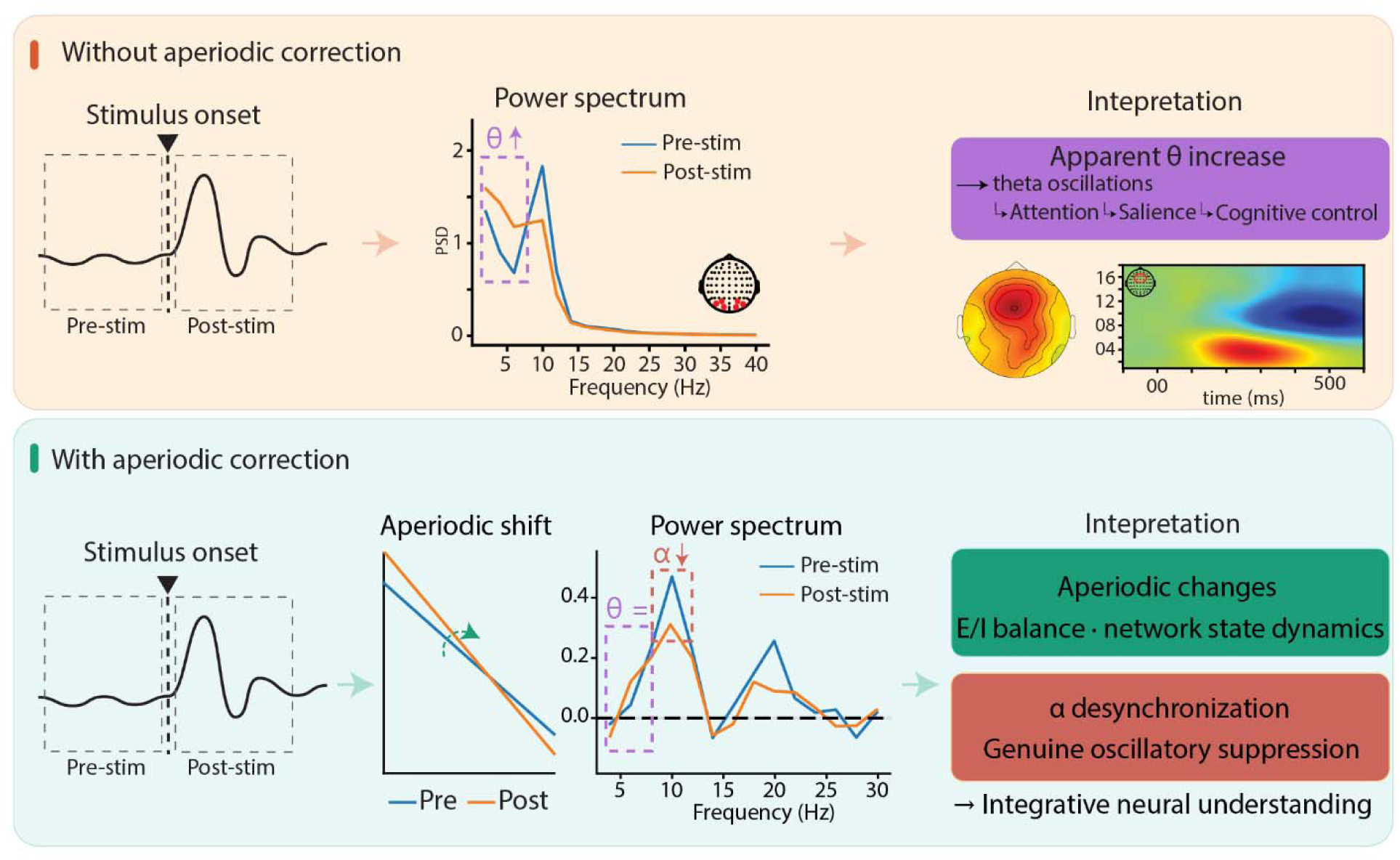

## Introduction

Neural activity in the frequency domain is shaped by at least two coexisting components: narrowband oscillatory rhythms and broadband, arrhythmic activity (Pritchard, 1992). Since the earliest days of EEG research (Berger, 1929), oscillations have dominated the interpretation of electrophysiological signals, leading to the canonical distinction between delta (< 4 Hz), theta (4-8 Hz), alpha (8-12 Hz), beta (12-30 Hz), and gamma (> 30 Hz) bands based on rhythms that emerge prominently across behavioral and cognitive states. This oscillation-centered framework has been highly productive and remains central to both basic and clinical neuroscience (Buzsaki & Draguhn, 2004). However, neural power spectra are not composed solely of rhythmic peaks. They also contain a broadband component, often described as scale-free or aperiodic activity, that follows a characteristic 1/f^x^-like profile, with power decreasing as frequency increases (Bullock et al., 1995; Freeman & Zhai, 2009; He, 2014; He et al., 2010; Miller, 2010). Thus, rather than reflecting only oscillatory processes, neural field signals such as EEG, MEG, and local field potentials are more accurately understood as the combination of rhythmic and arrhythmic activity.

This broadband aperiodic component was long treated as background noise and routinely removed to facilitate the study of oscillations (Cohen, 2014; Gyurkovics et al., 2021). More recently, however, growing evidence has suggested that aperiodic activity is itself functionally meaningful and may index ongoing changes in neural state (Donoghue et al., 2020; Gao et al., 2017; Manning et al., 2009; Miller et al., 2012; Preston et al., 2025; Waschke et al., 2021). In particular, the aperiodic exponent has been linked to the balance between excitatory and inhibitory synaptic activity, with steeper spectra thought to reflect relatively greater inhibition (Gao et al., 2017). Because of this, the shape of the aperiodic spectrum may provide a moment-to-moment marker of the underlying neural dynamics that support information processing.

Supporting this idea, recent work has shown that the aperiodic component changes following auditory stimulation and that these changes vary as a function of attentional context, such as whether a stimulus is frequent or rare (Gyurkovics et al., 2022). In that study, post-event spectra differed significantly from pre-event spectra even after removing the event-related time locked response. A substantial portion of this difference could be explained by a clockwise rotational shift in the 1/f component, characterized by increased low-frequency and decreased high-frequency power. Importantly, the magnitude of this rotational shift scaled with attentional demands, suggesting that stimulus processing is associated with systematic changes in aperiodic activity, potentially reflecting increased inhibition following stimulus onset.

In classic event-related time-frequency analyses, post-stimulus changes are made salient through some form of baseline correction where pre-stimulus activity is removed from the full epoch of interest (Gyurkovics et al., 2021). Using this framework, numerous studies have reported transient increases in theta power across a wide range of tasks and stimulus contexts (Cavanagh & Frank, 2014; Cavanagh & Shackman, 2015; Cavanagh et al., 2012; Cohen & Donner, 2013; Gyurkovics & Levita, 2021; McLoughlin et al., 2022; Vanneau, Quiquempoix, et al., 2025). More broadly, a substantial literature links frontal or midfrontal theta increases following a stimulus or event to control-related cognitive processes, although the precise functional interpretation varies across paradigms. As a general characterization, frontal-midline theta has often been viewed as a marker of control demand, attentional allocation, performance monitoring, or decision-related control, rather than of any single cognitive operation (Cavanagh & Frank, 2014). For example, theta activity has been reported to increase with cognitive load during working memory tasks (Hsieh & Ranganath, 2014; Jensen & Tesche, 2002). However, recent work in the context of working memory suggests that this well-known theta increase may be largely artifactual, arising from concurrent changes in aperiodic offset and exponent during the retention period rather than from a genuine increase in periodic theta activity (Frelih et al., 2025; van Engen et al., 2026).

While it is unambiguously clear that theta oscillations are a prominent feature of rodent hippocampus (Buzsaki et al., 2003), it remains unclear whether transient theta power increases reported in human cortical recordings in other contexts truly reflect oscillatory theta activity, or whether they are instead largely driven by changes in aperiodic activity. To address this question, we reanalyzed high-density EEG recorded in typically developing children (8-13 years old) data from two tasks. The first was an audiovisual simple reaction-time task involving stimuli from different sensory modalities, allowing us to test whether changes in aperiodic activity vary systematically as a function of sensory input (Sakowitz et al., 2000; Vanneau, Foxe, et al., 2025; Vicentin et al., 2024). The second was a classical visual oddball task. In this paradigm, many studies have reported enhanced transient fronto-central theta activity in response to rare target stimuli relative to frequent standard stimuli (Bernat et al., 2007; Cavanagh et al., 2012; Harper et al., 2019; Karakas et al., 2000), consistent with the view that frontal-midline theta indexes cognitive effort and top-down control, with greater theta associated with enhanced attention and performance. By comparing conventional spectral analyses with analyses that explicitly model and remove the aperiodic component, we sought to determine whether these apparent theta effects reflect genuine periodic activity or are instead largely attributable to changes in the aperiodic background.

## Methods

### Participants

Data collected for a different project were reanalyzed for the current study. The study initially included 41 typically developing children between 8 and 14 years of age. After excluding participants based on eye-tracking and EEG quality (see respective methods section for more details) the final analysis was conducted on 36 (10.43 years old ± 1.8; Mean ± Std) participants for the audiovisual task and 38 (10.44 years old ± 1.8) for the visual oddball task. Participants met the following inclusion criteria: no history of neurological, developmental, or psychiatric disorders, no first-degree relatives diagnosed with autism, and enrollment in age-appropriate grade in school. All procedures were approved by the Institutional Review Board of the Albert Einstein College of Medicine and adhered to the ethical standards outlined in the Declaration of Helsinki. All participants assented to the procedures, and their parents or guardian signed an informed consent approved by the Institutional Review Board of the Albert Einstein College of Medicine. Participants received nominal recompense for their participation (at $15 per hour).

### Audiovisual task: Experimental procedure (n = 36)

Participants were seated in a chair in an electrically shielded room (International Acoustics Company, Bronx, New York), 70 cm away from the visual display (Dell UltraSharp 1704FPT). The stimuli, controlled by Presentation software (Neurobehavioral Systems), included three types: a red disc (’Visual’), a 1000Hz tone (’Audio’), and their simultaneous presentation (’Audiovisual’). Participants were instructed to press a button as quickly as possible upon detecting any stimulus. The auditory stimulus was 1000Hz, 60ms tone presented binaurally (75 dB SPL). The visual stimulus was a red disc subtending to 1.5 degrees, displayed above a fixation cross. The audiovisual stimulus was a simultaneous presentation of both. Each trial presented a pseudo randomly chosen stimulus (A, V, or AV; represented equiprobably), with stimuli delivered through headphones (HD 650 Sennheiser) and displayed on a flat-panel LCD (Dell UltraSharp 1704FPT, 60Hz). A jittered randomly sampled interstimulus interval (1000-3000ms) reduced onset predictability and therefore anticipatory responses in the baseline period. The task consisted of 400 trials across 4 blocks (100 trials per block), each block lasting approximately 3 minutes and 40 seconds. Button presses were recorded using a response pad (Logitech Wingman Precision Gamepad). Triggers indicating stimulus onset were sent to the EEG stimulus channel from the PC acquisition computer via Presentation software.

### Visual Oddball task: Experimental procedure (n = 38)

Participants were seated in a chair in an electrically shielded room (International Acoustics Company, Bronx, New York), 70 cm away from the visual display (Dell UltraSharp 1704FPT). The stimuli, controlled by Presentation software (Neurobehavioral Systems), were objects, each shown as upright and inverted images, along with infrequently presented shadow versions (see Figure 4A). The objects come from the BOSS database (Brodeur et al., 2010). Participants were instructed to press a button as quickly as possible upon detecting a shadow stimulus (presented at 20% probability, see Figure 4A for illustration). A jittered interstimulus interval (900–1100ms) reduced onset predictability. The task comprised 360 trials across 6 blocks (60 trials/block, ∼1 minute per block).

### EEG recordings & preprocessing

EEG data were recorded at a sampling rate of 512Hz using 64 channels BioSemi Active II system (using the CMS/DRL referencing system) with an anti-aliasing filter (−3 dB at 3.6 kHz). Analyses were conducted in Python (3.11) using MNE(Gramfort et al., 2013) and custom scripts available at https://github.com/tvanneau/Ap-corrected. Bad channel detection was performed using the function NoisyChannels (with RANSAC) from the pyprep toolbox (Bigdely-Shamlo et al., 2015). If more than 15% of the channels were detected as bad, the participant was rejected. Bad channels were interpolated using spline interpolation (Perrin et al., 1989). EEG was filtered using a FIR band-pass filter (0.01-40Hz), and Independent Component Analysis (ICA) on 1Hz high-pass EEG was used to identify and manually reject eye-related components (blinks/saccades). For both tasks, epochs were created from −500 to +800ms around each stimulation with a baseline-correction from −50ms to +20ms stimulus onset and referenced to a common average reference.

### Spectral analysis pipeline

The same analysis pipeline was applied to both tasks. For each epoch, we defined a pre-stimulus window from −500 to 0ms and a post-stimulus window from 0 to 500ms. Power spectral density (PSD) was estimated separately within each window using Welch’s method with a Hann taper and constant detrending. Spectra were computed on a trial-by-trial basis for each EEG channel in the 2–35Hz range and then averaged across trials within each participant and condition. Results were robust to the choice of frequency range, as analyses conducted over 3–40 Hz yielded comparable findings. Frequency-band measures were subsequently derived for theta (4–8Hz), alpha (8–13Hz), beta (13–25Hz), and low gamma (25–35Hz). All analyses were performed on both the total signal and on ERP-removed data. For the latter, the condition-specific evoked response was first computed by averaging trials in the time domain and subtracted from each trial, thereby removing ERPs that have a broadband representation in the frequency domain and could thus have confounded aperiodic estimates (Gyurkovics et al., 2022), and the same PSD pipeline was then applied to these ERP-removed epochs. To compare conventional and aperiodic-aware estimates of stimulus-related spectral change, we implemented two parallel approaches. In the conventional approach, pre- and post-stimulus spectra were log-transformed and spectral change was quantified as the difference between post- and pre-stimulus PSD, i.e., log10(PSD_post_) − log10(PSD_pre_). Band-limited values were then obtained by averaging this spectrum within each frequency band of interest. In the aperiodic-corrected approach, the aperiodic component of both the pre- and post-stimulus spectra was parameterized using the specparam toolbox in fixed aperiodic mode. Model fitting was restricted to the 2–35Hz range. For each window, the fitted aperiodic component was subtracted from the log-transformed PSD to obtain an aperiodic-corrected residual spectrum. Spectral change was then quantified as the difference between the corrected post- and pre-stimulus spectra, i.e., [log10(PSD_post_) − Aperiodic_post_] − [log10(PSD_pre_) − Aperiodic_pre_]. Theta, alpha, beta, and gamma values were extracted from this residual change spectrum in the same way as for the conventional approach. In addition, the aperiodic parameters themselves were retained for both pre- and post-stimulus windows at each channel, including offset, exponent, model goodness-of-fit (R²), and fitting error.

### Statistical analysis

All statistical analyses were conducted using Jamovi (The jamovi project, version 2.3.28, https://www.jamovi.org) for linear mixed-effects models (LME) and Python (MNE-Python) for spatial permutation-based testing. Changes in band-limited spectral activity were analyzed using LME in order to account for repeated measurements within individuals. The fixed factors were condition (auditory, visual, audiovisual for the audiovisual task; standard, target for the oddball task) and method (conventional, aperiodic-corrected). Participant was included as a clustering variable with a random intercept, capturing between-participant variability in baseline responses. Fixed effects were evaluated using Type III Wald F-tests (Fixed Effects Omnibus Tests), which test each fixed effect at the parameter estimates of the fitted model while holding all other effects constant. This approach does not involve sequential comparison of nested models; rather, it provides an omnibus test of each effect within the single fitted model, analogous in logic to a conventional ANOVA but conducted within the mixed-effects framework. For each model, we report the omnibus F statistic, the associated degrees of freedom, and the corresponding p value. As a standardised measure of effect size, we report partial eta squared (η²p), computed from the F statistic and its associated degrees of freedom as η²p = (F × df_effect) / (F × df_effect + df_error), which provides an approximation of the proportion of variance in the outcome attributable to each fixed effect after accounting for other terms in the model. Statistical significance was set at α = 0.05. Residual normality was evaluated using the Shapiro–Wilk test. To identify brain regions showing significant changes in aperiodic parameters (offset and exponent), we used a nonparametric spatial cluster-based permutation test (Maris & Oostenveld, 2007) as implemented in MNE-Python. For each condition, we first computed subject-level change values at each electrode as the difference between post- and pre-stimulus estimates. These channel-wise change scores were then entered into a one-sample cluster-based permutation test against zero across participants. Clusters were formed over spatially adjacent electrodes using the BioSemi64 sensor adjacency matrix, with a two-sided cluster-forming threshold corresponding to α = 0.01. Cluster significance was assessed using 5000 permutations based on random sign-flipping of the subject-level change values under the null hypothesis of no systematic pre–post difference. For each permutation, the maximum cluster-level statistic was retained to construct the null distribution, and observed clusters were assigned p-values by comparison with this distribution. In this implementation, the cluster-level statistic corresponded to cluster mass, that is, the sum of t-values within each spatial cluster (Maris & Oostenveld, 2007).

## Results

### Stimulus-evoked changes in aperiodic activity are modality-specific and persist after ERP removal

Examining aperiodic activity before and after stimulus onset revealed consistent findings across both paradigms. In the multisensory task, all sensory conditions were associated with a stimulus-related increase in the aperiodic exponent, indicating a steepening of the 1/f component following stimulation. Although this increase was present across conditions, its topographical distribution was modality-specific (Fig. 2B): auditory stimulation produced a predominantly central pattern, visual stimulation a more parieto-occipital pattern, and audiovisual stimulation a combination of both. Similar results were observed for the aperiodic offset. In the visual oddball task, a significant post-stimulus increase in both the aperiodic exponent and offset was observed for both standard and target trials, with significant clusters spanning all channels for the total signal and a large central region after ERP removal (Fig. 4B), indicating a stimulus-induced steepening of the aperiodic slope accompanied by a positive shift in offset regardless of attentional demands.

**Figure 1.**
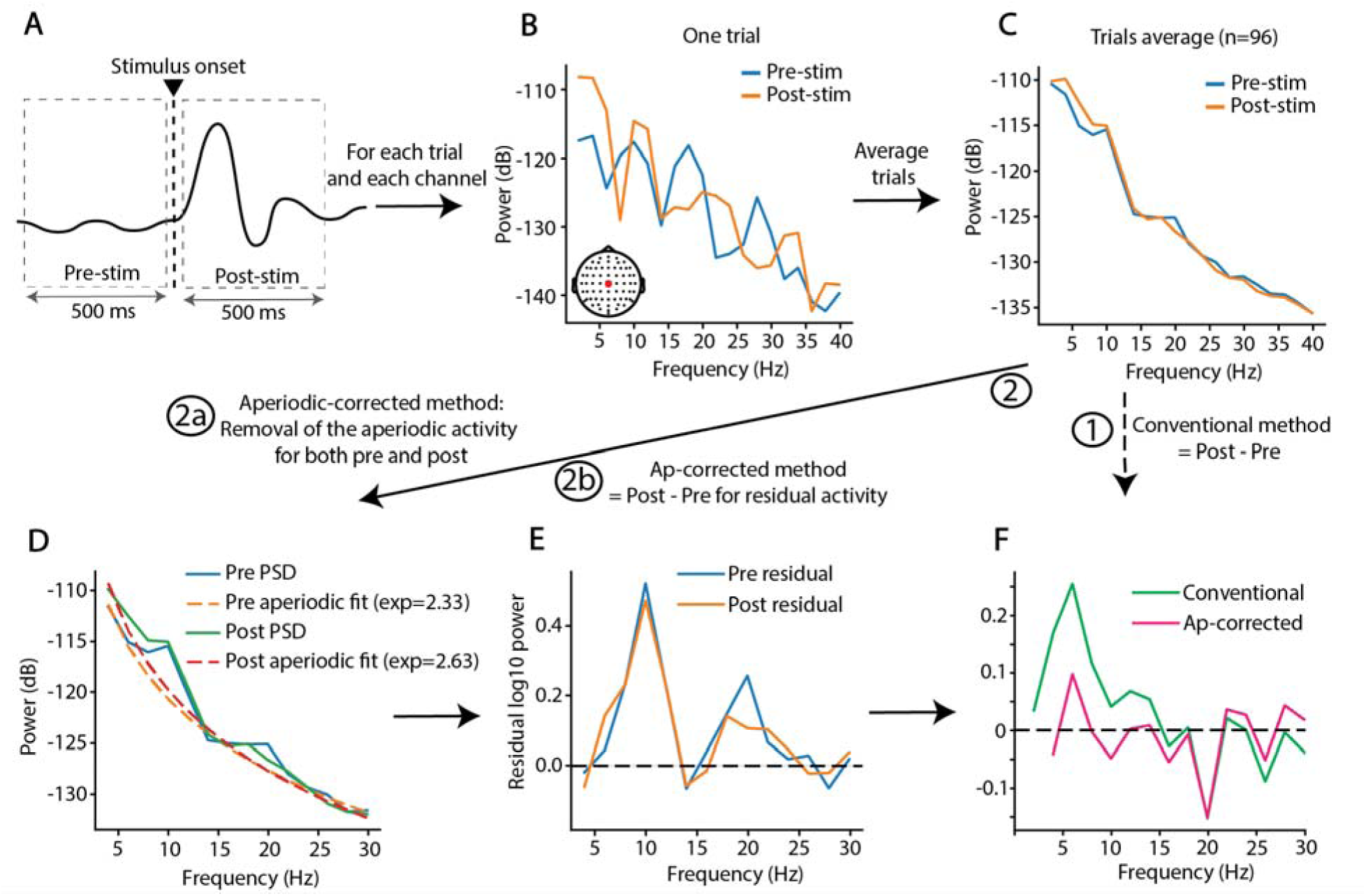
Spectral analysis pipeline. **(A)** For each trial, pre-stimulus (−500 to 0ms) and post-stimulus (0 to 500ms) windows were extracted. **(B)** Power spectral density (PSD) was computed separately for each window using Welch’s method with a Hann taper and constant detrending. **(C)** Trial-wise PSDs were averaged within each stimulus condition. From these spectra, two complementary approaches were used to quantify stimulus-related spectral change: **(1)** a conventional approach based on log-transformed pre/post spectral change, and **(2)** an aperiodic-corrected approach. **(D)** The aperiodic component of the pre- and post-stimulus spectra was parameterized independently using the specparam toolbox (Donoghue et al., 2020). **(E)** Aperiodic-corrected residual spectra were obtained by subtracting the fitted aperiodic component from each window-specific PSD. **(F)** Spectral change was then quantified as the post-minus-pre difference, either from the uncorrected spectra (conventional method, green) or from the aperiodic-corrected residual spectra (aperiodic-corrected method, magenta).

**Figure 2.**
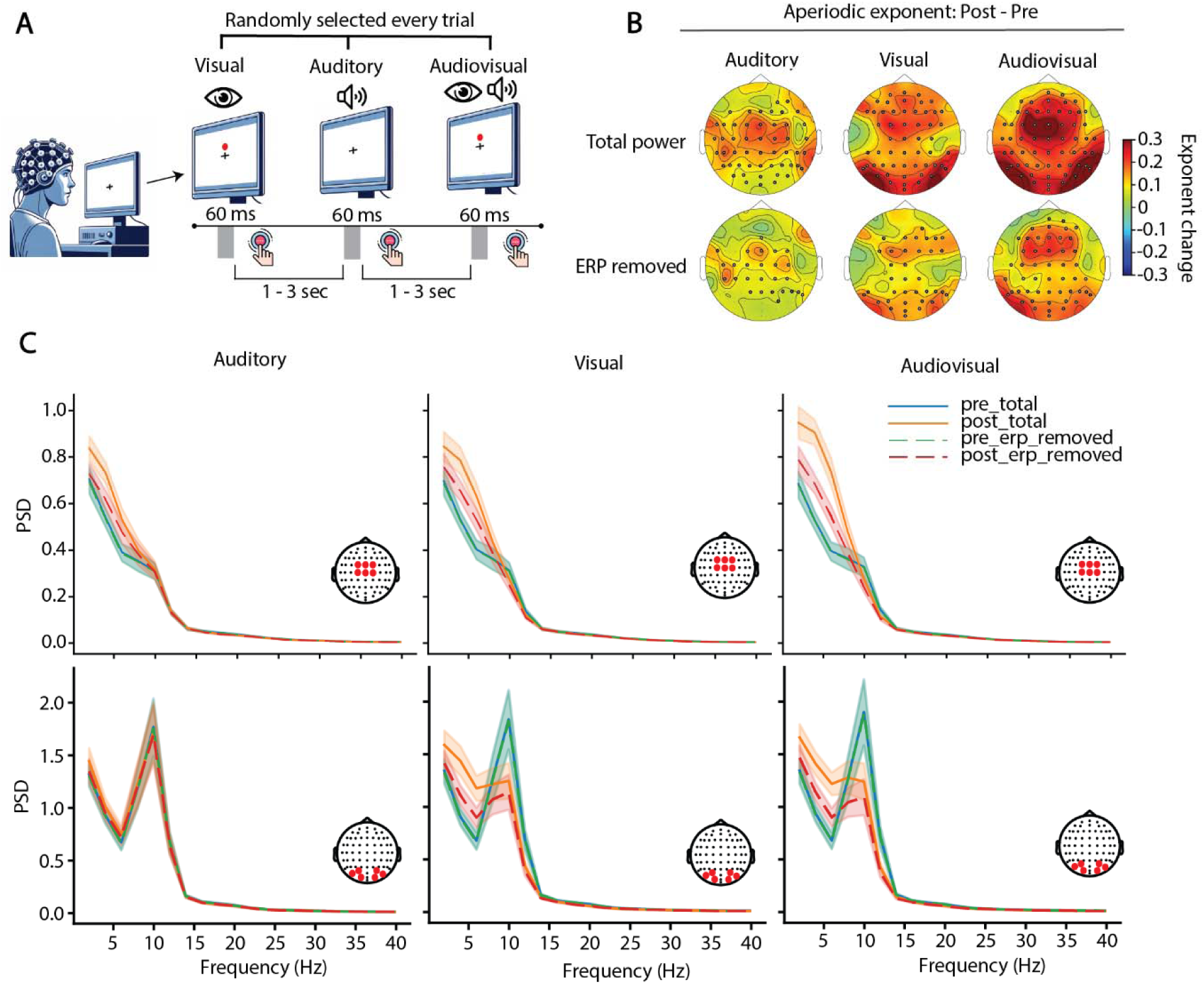
Aperiodic activity changes following stimulus onset and varies as a function of sensory modality. **(A)** Schematic of the audiovisual simple reaction-time task. Visual-only, auditory-only, and audiovisual stimuli were presented in random order, and participants were instructed to respond to every stimulus. **(B)** Topographical maps of the change in aperiodic exponent from pre-to post-stimulus window (post − pre) for each stimulus condition. Results are shown for the total signal (top row) and after time-domain removal of the ERP (bottom row). Black dots indicate channels belonging to significant spatial clusters identified using cluster-based permutation testing against zero change. **(C)** Power spectral density (PSD, linear domain; see Supplementary figure 1 for PSD in log) in the pre-stimulus window (blue) and post-stimulus window (orange), shown for the total signal (solid lines) and after time-domain ERP removal (dashed lines), for auditory (left), visual (middle), and audiovisual (right) stimulation. Spectra are averaged across a fronto-central channel cluster (top row) and a parieto-occipital channel cluster (bottom row).

Across both paradigms, time-domain ERP removal reduced the magnitude of these aperiodic effects but did not eliminate them: significant spatial clusters remained and, in the multisensory task, preserved the same overall modality-specific topographical organization (Fig. 2B). This was further supported by PSDs extracted from central and parieto-occipital channel clusters for each sensory condition, where post-stimulus spectra showed clear changes in offset and exponent relative to the pre-stimulus window that remained visible after ERP removal, albeit attenuated (Fig. 2C). Together, these findings indicate that the ERP contributes to stimulus-related aperiodic changes but does not fully account for them. Importantly, these broadband spectral shifts reshaped the spectral profile in ways that generate an apparent increase in theta-range power and reduce the observed magnitude of alpha desynchronization.

### Theta-band: Aperiodic correction abolishes the apparent stimulus-related increase

Prior to aperiodic correction, theta-band power changes displayed a topographical pattern closely mirroring aperiodic effects. In the multisensory task, auditory stimulation was associated with increased theta-band power over central channels, visual stimulation with increases over both central and parieto-occipital channels, and audiovisual stimulation with a similar but spatially broader pattern (Fig. 3A). Following aperiodic correction, the theta-band increase was greatly reduced for all stimulus types. In the visual oddball task, a similar pattern was observed, with fronto-central theta increases for both standard and target stimuli that were substantially reduced after aperiodic correction (Fig. 4A).

**Figure 3.**
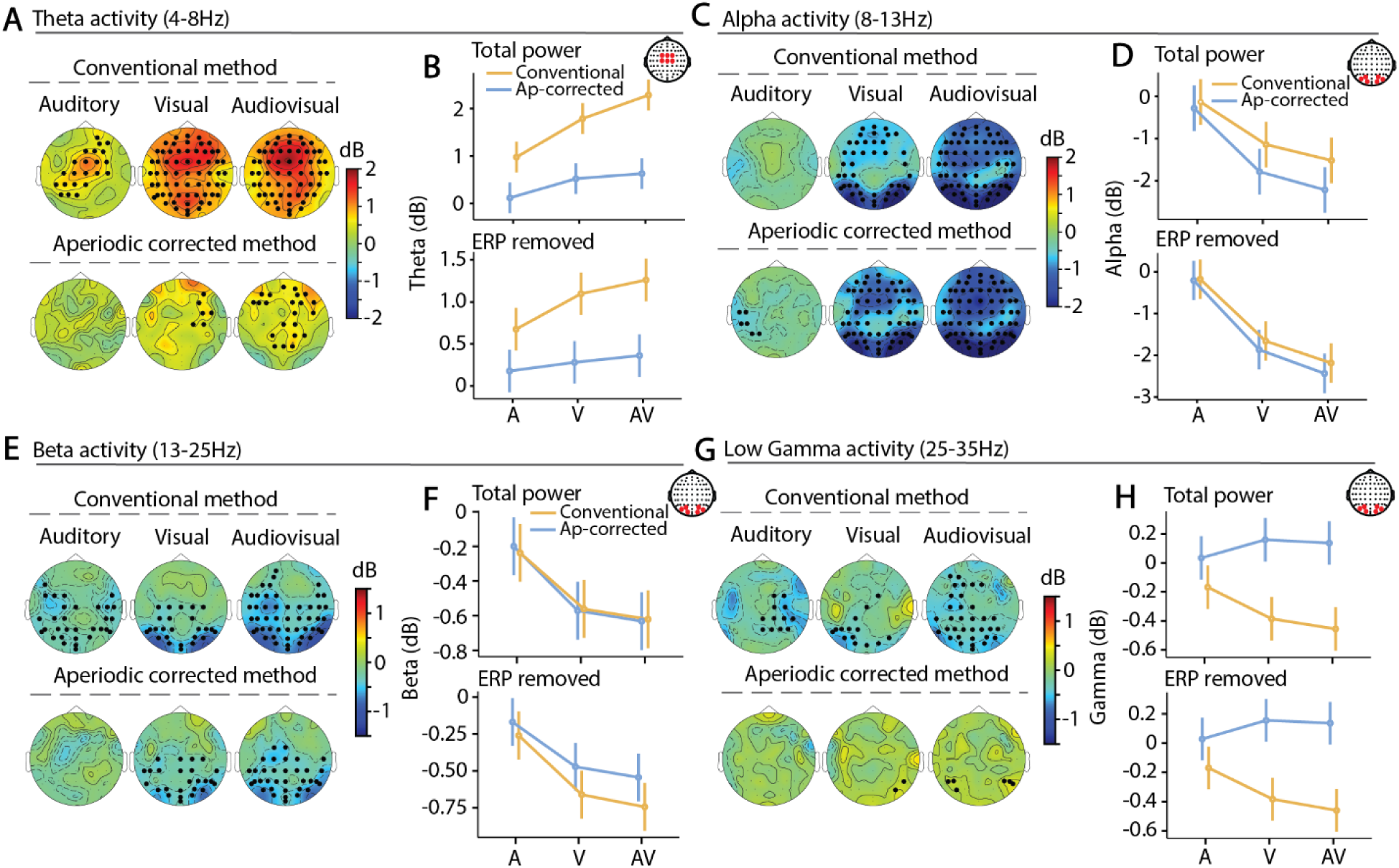
Aperiodic shifts explain apparent theta and gamma changes but not alpha and beta desynchronization across sensory modalities. **(A)** Topographical representation of theta-band (4–8 Hz) changes in response to auditory (left), visual (middle), and audiovisual (right) stimulation after removal of the ERP. Theta was quantified either with the conventional approach (top row; post–pre change in log-transformed PSD) or after aperiodic correction of both pre- and post-stimulus spectra (bottom row; see Methods). **(B)** Theta-band values averaged across a fronto-central channel cluster for each stimulus condition, shown for the conventional method (orange) and the aperiodic-corrected method (blue), for the total signal (top panel) and ERP-removed signal (bottom panel). **(C**–**D)** Same as **(A**–**B)** for alpha-band activity (8–13 Hz), **(E-F)** for beta-band activity (13-25Hz), **(G-H)** for low gamma-band activity (25-35Hz).

**Figure 4.**
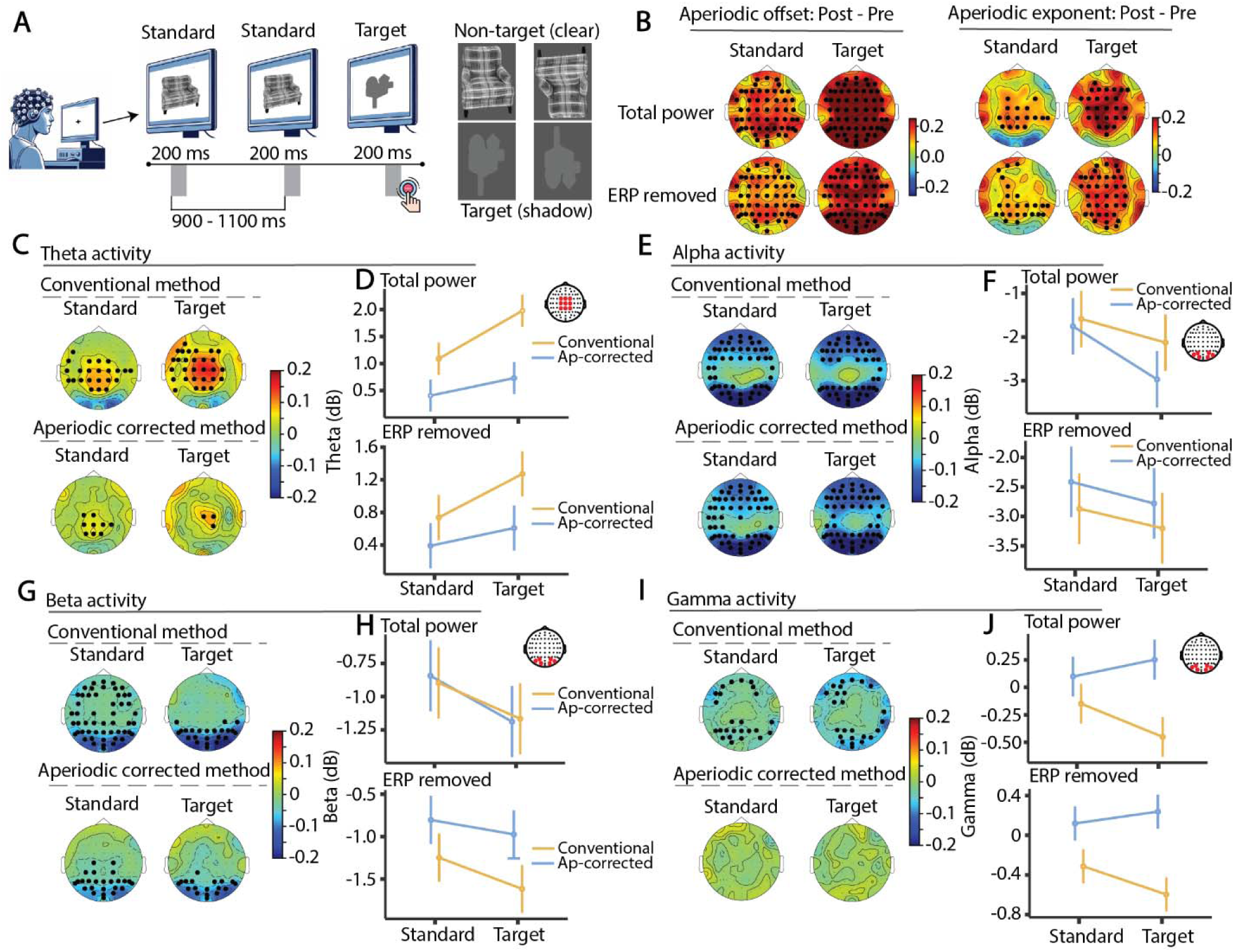
Aperiodic activity accounts for most stimulus-related theta changes, but not alpha changes, during the visual oddball task. **(A)** Schematic of the visual oddball task. Participants were presented with upright and inverted clear object stimuli (standards) and shadow versions (targets) and were instructed to respond only to target stimuli. **(B)** Topographical maps of the change in aperiodic exponent from pre-to post-stimulus window (post − pre) for standard and target stimuli, shown for the total signal (left) and after time-domain ERP removal (right). Black dots indicate channels belonging to significant spatial clusters identified with cluster-based permutation testing against zero change. **(C)** Topographical maps of theta-band (4–8 Hz) changes in response to standard (left) and target (right) stimuli, computed either with the conventional method (top row; post–pre change in log-transformed PSD) or after aperiodic correction of both pre- and post-stimulus spectra (bottom row; see Methods). **(D)** Mean theta-band values averaged across a fronto-central channel cluster for standard and target stimuli, shown for the conventional method (orange) and the aperiodic-corrected method (blue), for the total signal (top panel) and ERP-removed signal (bottom panel). **(E**–**F)** Same as **(C**–**D)** for alpha-band activity (8–13 Hz), **(G-H)** for beta-band activity (13-25Hz) and **(I-J)** for low gamma-band activity (25-35Hz).

**Figure 5.**
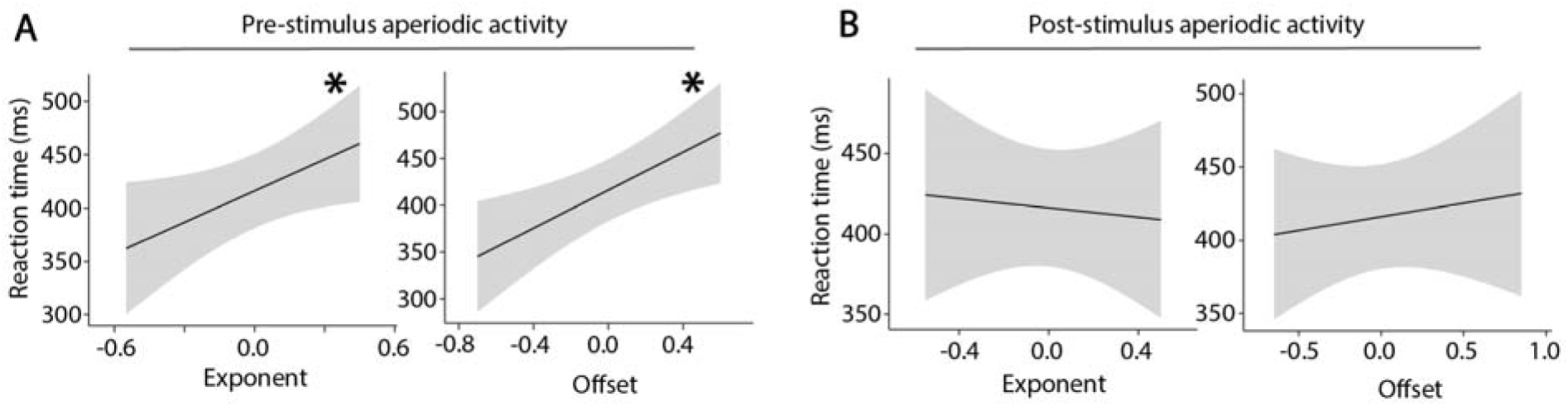
Relationship between aperiodic activity and reaction time. (**A**) Linear mixed-effects model predictions showing the association between pre-stimulus aperiodic exponent (left) and offset (right) and reaction time, while controlling for sensory modality (condition as a fixed effect) and including Subject as a random intercept. Aperiodic parameters are mean-centered such that zero corresponds to the sample mean. (**B**) Same analysis for post-stimulus averaged aperiodic parameters.

To formally test whether each method yielded statistically significant pre-to post-stimulus changes in theta-band power, we conducted spatial cluster-based permutation tests separately for each method and each sensory condition. In the multisensory task, the conventional method yielded a significant spatial cluster confined to central channels for auditory stimulation, and clusters encompassing a much broader scalp distribution for both visual and audiovisual stimulation. After aperiodic correction, no significant theta-band cluster was found for auditory stimulation, and only small, spatially restricted clusters emerged for visual (right frontal) and audiovisual (frontal and parietal) stimulation, indicating that the pre-to post-stimulus theta-band change is substantially reduced and largely non-significant once aperiodic activity is accounted for (Fig. 3A). In the oddball task, both standard and target stimuli yielded significant fronto-central theta clusters for the conventional method; after aperiodic correction, no significant clusters were observed for either condition (Fig. 4A).

Given that the largest theta-band changes were consistently observed over fronto-central channels, and that this region showed maximal effect amplitudes across conditions, theta-band power changes were averaged across a fronto-central ROI (Cz, FC1, FCz, FC2, Fz) and submitted to linear mixed-effects models with stimulus condition and method (conventional vs. aperiodic-corrected) as fixed factors and subject as a random factor. Models were conducted separately for the total signal and the ERP-removed signal.

In the multisensory task and for the total signal, there was a significant main effect of condition (F(2,175) = 36.73, η²p = 0.30, p < 0.001), with theta-band changes increasing progressively from auditory to visual to audiovisual stimulation, and significant pairwise differences between all three conditions (all ps ≤ 0.017). There was also a robust main effect of method (F(1,175) = 203.35, η²p = 0.54, p < 0.001), with substantially smaller theta-band changes after aperiodic correction (Δ = 1.28 dB ± 0.08, t = 14.3). Critically, the condition × method interaction was significant (F(2,175) = 6.76, η²p = 0.07, p < 0.001): whereas the conventional method showed clear differences across all stimulus conditions, these differences were markedly reduced after aperiodic correction, with only the auditory versus audiovisual comparison remaining significant and with a much smaller effect size (conventional: t = 8.55, p < 0.001; aperiodic-corrected: t = 3.35, p = 0.015; V vs AV for conventional: t = 3.25, p = 0.02; aperiodic-corrected: t = 0.7, p = 1.0). The same general pattern was observed after ERP removal, with theta-band changes remaining significantly larger for the conventional method (F(1,175) = 96.31, η²p = 0.35, p < 0.001), though the condition × method interaction did not reach significance (F(2,175) = 2.66, η²p = 0.03, p = 0.072).

A strikingly similar pattern emerged in the visual oddball task. For the total signal, there was a significant main effect of condition (F(1,111) = 24.43, η²p = 0.18, p < 0.001), with greater theta-band increases for targets than for standards (Δ = 0.6 dB ± 0.1, t = 4.94, p < 0.001), and a significant main effect of method (F(1,111) = 61.71, η²p = 0.36, p < 0.001), reflecting larger theta-band changes in the conventional analysis (Δ = 1.0 dB ± 0.1, t = 7.86, p < 0.001). The condition × method interaction was significant (F(1,111) = 5.27, η²p = 0.05, p = 0.02): the target-related theta enhancement was present with the conventional method (Δ = 0.9 dB ± 0.1, t = 5.12, p < 0.001) but was no longer significant after aperiodic correction (Δ = 0.3 dB ± 0.1, t = 1.87, p = 0.38). After ERP removal, the main effects of condition and method remained significant (ps < 0.001), and although the interaction did not reach significance (F(1,111) = 2.03, η²p = 0.01, p = 0.15), the same overall pattern was observed: target-related theta enhancement remained significant for the conventional method (p = 0.005) but not for the aperiodic-corrected method (p = 0.99).

As an exploratory analysis, we contrasted theta-band changes before and after ERP removal separately for each method and task. In both paradigms, ERP removal was associated with a significant reduction in theta-band change magnitude for the conventional method (multisensory: F(1,175) = 74.34, η²p = 0.3, p < 0.001; oddball: F(1,111) = 20.31, p < 0.001), but this effect was absent or negligible for the aperiodic-corrected method (multisensory: F(1,175) = 3.93, η²p = 0.02, p = 0.049; oddball: F(1,111) < 0.1, p = 0.93). This indicates that the conventional method is considerably more sensitive to ERP contamination than the aperiodic-corrected method, and that a large proportion of the theta-band change it detects reflects the ERP rather than oscillatory dynamics.

Together, and across both paradigms, these results indicate that the apparent stimulus-related theta-band increase is largely explained by concurrent changes in aperiodic activity rather than by a genuine increase in oscillatory theta power.

### Alpha-band: Effects remain present even after aperiodic correction

Unlike theta, alpha-band activity was characterized by a decrease in power following stimulus onset, consistent with event-related alpha desynchronization. In the multisensory task, auditory stimulation produced only modest desynchronization primarily over temporal channels, whereas visual and audiovisual stimulation elicited a much stronger desynchronization over parieto-occipital regions (Fig. 3C). In the oddball task, the spatial distribution was very similar between standard and target stimuli and between methods, with significant parieto-occipital and frontal clusters identified for all conditions (Fig. 4E). Notably, in both paradigms, aperiodic correction modulated the magnitude of alpha desynchronization, consistent with the idea that stimulus-related changes in aperiodic activity partially mask the true extent of alpha suppression.

At the cluster level in the multisensory task, significant alpha-band parieto-occipital clusters were identified by both methods for visual and audiovisual stimulation, but the aperiodic-corrected method additionally revealed a significant temporal cluster absent from the conventional analysis and produced a spatially broader central cluster for visual stimulation, a pattern that stands in clear contrast to the theta-band findings. In the oddball task, significant clusters were identified by both methods for standard and target stimulation across all conditions.

Alpha changes were averaged across parieto-occipital ROI (O1, Oz, O2, PO3, PO7, PO4 and PO8) and submitted to linear mixed-effects models. In the multisensory task and for the total signal, there was a significant main effect of condition (F(2,175) = 49.42, η²p = 0.36, p < 0.001), with stronger alpha desynchronization for visual and audiovisual stimuli than for auditory stimuli (A vs V: Δ = 1.25 dB, t = 7.22, p < 0.001; A vs AV: Δ = 1.66 dB, t = 9.53, p < 0.001), and a significant main effect of method (F(1,175) = 12.17, η²p = 0.07, p < 0.001), reflecting greater desynchronization after aperiodic correction (Δ = 0.5 dB, t = 3.49, p < 0.001), but no significant condition × method interaction (F(2,175) = 1.51, η²p = 0.02, p = 0.22). In the oddball task, there was similarly a significant main effect of condition (F(1,111) = 40.04, η²p = 0.26, p = 0.005), with stronger alpha desynchronization for target than standard stimuli, a significant main effect of method (F(1,111) = 13.18, η²p = 0.1, p < 0.001), and a significant condition × method interaction (F(1,111) = 5.91, η²p = 0.05, p = 0.01), with a larger standard-target difference for the aperiodic-corrected method (Δ = 1.2 dB, t = 6.19, p < 0.001) than for the conventional method (Δ = 0.5 dB, t = 2.75, p = 0.04).

After ERP removal, the pattern diverged somewhat between paradigms. In the multisensory task, the condition effect remained significant (F(2,175) = 102.34, η²p = 0.54, p < 0.001), while neither the main effect of method nor the condition × method interaction reached significance (ps > 0.12). In the oddball task, the condition effect again remained significant (F(1,111) = 8.31, η²p = 0.07, p = 0.005), but in contrast to the multisensory task, a significant main effect of method emerged with smaller alpha desynchronization for the aperiodic-corrected method (F(1,111) = 13.15, η²p = 0.1, p < 0.001), with no significant interaction.

Contrasting alpha changes before and after ERP removal revealed a consistent pattern across paradigms: ERP removal was associated with a significant increase in alpha desynchronization for the conventional method (multisensory: F(1,175) = 8.11, p = 0.005; oddball: F(1,111) = 96.52, p < 0.001), indicating that the ERP attenuates the underlying alpha desynchronization signal with this approach. No such effect was observed for the aperiodic-corrected method in either paradigm (multisensory: F(1,175) = 0.37, p = 0.54; oddball: F(1,111) = 2.4, p = 0.12).

Together, these results suggest that alpha desynchronization is not primarily driven by changes in aperiodic activity and remains robust after aperiodic correction, in clear contrast to the theta-band findings. Aperiodic correction modulates the estimated magnitude of alpha suppression, likely by removing the attenuating influence of concurrent broadband shifts but does not abolish the effect.

### Beta-band: Aperiodic correction reduces the apparent magnitude of desynchronization

Beta-band activity exhibited a post-stimulus decrease in power relative to baseline in both paradigms (Figs. 3E, 4E). In the multisensory task, significant parieto-occipital beta-band clusters were identified by both methods for visual and audiovisual stimulation, though these were spatially broader for the conventional method, and no significant cluster was observed for auditory stimulation after aperiodic correction. A similar pattern was observed in the oddball task, where both methods yielded significant parieto-occipital clusters for standard and target stimulation, again with broader spatial extent for the conventional method.

Beta-band power was averaged across the parieto-occipital ROI used for alpha and submitted to linear mixed-effects models. When considering the total signal, results were comparable across both paradigms: only the main effect of condition reached significance (multisensory: F(2,175) = 21.68, η²p = 0.20, p < 0.001; oddball: F(1,111) = 13.89, η²p = 0.11, p < 0.001), with no significant effect of method or condition × method interaction (all ps > 0.6). After ERP removal, the condition effect remained significant in both paradigms (multisensory: F(2,175) = 26.16, η²p = 0.23, p < 0.001; oddball: F(1,111) = 14.79, η²p = 0.12, p < 0.001), and a significant main effect of method emerged in both, indicating that the conventional method overestimated the magnitude of beta desynchronization relative to the aperiodic-corrected method (multisensory: Δ = 0.16 dB, t = 3.1, p = 0.002; oddball: Δ = 0.5 dB, t = 7.8, p < 0.001). The condition × method interaction remained non-significant in both paradigms.

Contrasting beta changes before and after ERP removal revealed different patterns for the two methods. For the conventional method, ERP removal was associated with a significant change in beta desynchronization magnitude in the oddball task (F(1,111) = 51.57, p < 0.001) but not in the multisensory task (F(1,175) = 2.86, p = 0.09). The aperiodic-corrected method showed no significant effect of ERP removal in either paradigm (multisensory: F(1,175) = 2.51, p = 0.11; oddball: F(1,111) = 3.61, p = 0.06), suggesting that ERP contamination does not primarily account for the discrepancy between methods in this frequency band.

Together, these results indicate that while the two methods yield comparable estimates of beta desynchronization for the total signal, the conventional method systematically overestimates the magnitude of beta desynchronization in the ERP-removed signal, consistent with a contribution of aperiodic slope changes to broadband power that disproportionately affects the conventional approach.

### Gamma-band: Aperiodic correction reverses the apparent direction of activity changes

Finally, we examined stimulus-related changes in the low gamma band (25–35 Hz), where the two approaches yielded qualitatively opposite patterns in both paradigms. In the multisensory task, the conventional method identified significant gamma desynchronization clusters across all three conditions, whereas after aperiodic correction no desynchronization was observed; instead, significant gamma-band synchronization emerged, with a right temporal cluster for visual stimulation and bilateral temporal clusters for audiovisual stimulation (Fig. 3G). In the visual oddball task, the conventional method similarly identified significant gamma desynchronization clusters across parieto-occipital and frontal channels for both standard and target stimulation, while after aperiodic correction no significant changes in gamma activity were detected (Fig. 4I).

Linear mixed-effects models confirmed a highly significant effect of method in both paradigms (multisensory: F(1,175) = 58.74, η²p = 0.25, p < 0.001; oddball: F(1,111) = 32.72, η²p = 0.22, p < 0.001): while the conventional method indicated desynchronization, the aperiodic-corrected method indicated either synchronization (multisensory) or no change (oddball), with a consistent difference between methods of approximately 0.45–0.47 dB. The main effect of condition was not significant in either paradigm, but significant condition × method interactions were observed in both (multisensory: F(2,175) = 4.47, η²p = 0.05, p = 0.01; oddball: F(1,111) = 7.53, η²p = 0.06, p = 0.007), though no individual post-hoc comparisons reached significance in either case.

In contrast to the theta and alpha bands, contrasting gamma changes before and after ERP removal revealed no significant effect of signal type for either method in either paradigm (all ps > 0.09), indicating that ERP contamination does not account for the discrepancy between methods in this frequency band.

Taken together, these findings demonstrate that the apparent gamma-band desynchronization detected by the conventional method is a spurious consequence of concurrent changes in aperiodic activity. Specifically, the post-stimulus increases in both the exponent and offset of the aperiodic slope steepens the spectral gradient, artificially suppressing power at higher frequencies and producing the illusion of gamma desynchronization in the absence of any true change in narrowband gamma oscillatory activity.

### Pre-stimulus aperiodic activity is associated with reaction time

We next examined whether aperiodic activity was related to behavioral performance. For the audiovisual task, linear mixed-effects models were fitted with reaction time as the dependent variable, sensory modality as a fixed factor, and the aperiodic parameter of interest entered as a covariate. These analyses revealed that both pre-stimulus aperiodic parameters were significantly associated with reaction time, for both the exponent (F(1,102) = 4.27, η²p = 0.04,p = 0.041, β = 93.7ms) and the offset (F(1,99) = 8.67, η²p = 0.08, p = 0.004, β = 100.8ms). This is consistent with recent evidence suggesting that flatter pre-stimulus aperiodic slopes, reflecting broader cortical excitability, are associated with better perceptual performance (Cunningham et al., 2023). In contrast, post-stimulus aperiodic parameters were not significantly associated with reaction time, possibly reflecting increased spectral variability during the post-stimulus period that reduces the reliability of the aperiodic estimate as a behavioral predictor. In contrast, post-stimulus aperiodic parameters were not significantly associated with reaction time. For the visual oddball task, reaction times were available only for target stimuli. Because the number of target trials was lower than in the audiovisual task and only a single condition was considered, these results should be interpreted more cautiously. Nonetheless, a similar pattern was observed, with a significant association between pre-stimulus offset and reaction time (Supplementary Figure 5). Taken together these results suggest that flatter aperiodic activity before the stimulus is associated with faster reaction time.

## Discussion

The present study investigated event-related changes in the aperiodic, 1/f-like component of the EEG spectrum and its consequence on conventional baseline-normalized spectral analyses. Across two distinct paradigms, stimulus onset reliably elicited a steepening of the aperiodic exponent, with modality- and task-specific spatial distributions that persisted after ERP removal. These broadband spectral shifts were not inconsequential: because the aperiodic change took the form of a clockwise rotation of the power spectrum, it systematically distorted frequency-specific estimates of post-stimulus power changes across the full spectrum. At lower frequencies, this rotation artificially inflated power, producing spurious theta-band increases and partially masking oscillatory alpha desynchronization; at higher frequencies, it suppressed power, artificially amplifying apparent beta-band and gamma-band desynchronization. Consequently, conventional spectral analyses mischaracterized the direction and/or magnitude of stimulus-related changes across all examined frequency bands. These findings suggest that many previous reports of transient post-stimulus theta increases may have overestimated the contribution of genuine periodic oscillations by conflating them with shifts in the aperiodic background. More broadly, they call for greater caution when interpreting frequency-specific post-stimulus activity using conventional normalization approaches that implicitly assume aperiodic stationarity across the peri-stimulus window.

### Abolishment of apparent theta oscillatory activity when disentangling periodic and aperiodic components in EEG activity

This study adds to the growing literature on the limitations of baseline correction for spectral analysis, where pre-stimulus activity is removed from the full epoch of interest (Cohen, 2014; Gyurkovics et al., 2021). Decomposing the EEG signal into periodic and aperiodic components provides a more veridical picture of spectral dynamics (rather than relying on baseline normalization), revealing that low-frequency theta activity largely “disappears” from the periodic component once the aperiodic background is accounted for. These findings are consistent with a series of recent studies investigating aperiodic activity changes in working memory and cognitive control contexts (Akbarian et al., 2024; Frelih et al., 2025; Kalamala et al., 2024; Lendner et al., 2023; Preston et al., 2025; van Engen et al., 2026; Virtue-Griffiths et al., 2025), converging on the conclusion that most stimulus- or task-related theta changes in human EEG are accounted for by aperiodic modulation rather than by genuine oscillatory dynamics. Here we extend those findings by showing that aperiodic changes are modality-specific, localizing to sensory areas in response to each sensory modality, consistent with previous findings in an audiovisual detection task (Waschke et al., 2021), and that attentional demands further modulate aperiodic activity in the context of a visual oddball, paralleling similar observations in auditory oddball paradigms (Gyurkovics et al., 2022).

### Reduction of the apparent beta and gamma desynchronization after aperiodic correction

The same logic extends to the higher frequency bands examined here. In the beta band, the conventional method (subtraction of the full pre-stimulus spectrum from the full post-stimulus spectrum) systematically overestimated the magnitude of desynchronization, particularly in the ERP-removed signal, consistent with the aperiodic slope change suppressing power broadly at frequencies above the spectral rotation point. In the gamma band, the distortion was most striking: what appeared as gamma desynchronization under conventional analysis was entirely reversed after aperiodic correction, revealing either synchronization or no change, indicating that the apparent gamma suppression was a purely spurious consequence of spectral slope steepening. It is worth noting that the stimuli used in the present study would not be expected to elicit robust gamma increases of the kind typically reported in response to luminance gratings (Hermes et al., 2015). This finding should be interpreted with some caution, however, as gamma-band activity is inherently more difficult to measure reliably with scalp EEG due to its low signal-to-noise ratio and susceptibility to muscular artifacts (Muthukumaraswamy, 2013; Nunez & Srinivasan, 2010). Nonetheless, the consistency of this reversal across both paradigms suggests that in the case the apparent modulation of gamma activity reflects a genuine methodological artifact rather than random noise.

Together, these results suggest that the clockwise rotation of the power spectrum following stimulus onset produces a systematic pattern of distortion across frequency bands that, under conventional analyses, manifests as apparent theta and alpha increases at low frequencies and apparent desynchronization in the beta and gamma ranges, a comprehensive set of effects that span much of the frequency range typically analyzed in EEG research.

### Alpha desynchronization appears to reflect genuine periodic suppression

In contrast to the theta, beta, and gamma findings, alpha desynchronization was not diminished by aperiodic correction, it was enhanced. After accounting for the aperiodic component, the magnitude of alpha suppression was larger than estimated by the conventional method, and the effect remained robust across both paradigms and both the total and ERP-removed signals. This pattern is consistent with the interpretation that stimulus-related changes in the aperiodic slope partially mask the true extent of alpha desynchronization: the broadband upward shift in power caused by the clockwise spectral rotation attenuates the apparent dip in the alpha range, such that correcting for this shift unmasks a stronger suppression signal. Aperiodic correction also improved spatial sensitivity to alpha desynchronization, revealing additional temporal clusters in the multisensory task that were absent from the conventional analysis.

Although the oscillatory character of the signal in the time-domain was not measured directly (Karvat et al., 2026; Myrov et al., 2024), the fact that alpha desynchronization only increased in effect size after accounting for aperiodic dynamics suggests that the alpha decrease, unlike the apparent theta increase, reflects genuine oscillatory suppression rather than an artifact of aperiodic changes. This distinction is important for interpreting the functional significance of these effects. Alpha desynchronization is widely described as an index of cortical excitability (Klimesch et al., 2007): in response to both auditory and visual input, alpha power over the corresponding sensory cortex decreases and is associated with increased cortical excitability, faster reaction times, and more efficient sensory processing (Darrell et al., 2026; Lange et al., 2013; Romei et al., 2008; Samaha et al., 2017; Sauseng et al., 2009; Sauseng et al., 2005; Vanneau, Foxe, et al., 2025). The fact that aperiodic correction preserves and even amplifies this effect reinforces its status as a reliable index of sensory cortical dynamics, while at the same time revealing that conventional analyses have been systematically underestimating its magnitude.

One intriguing question raised by these findings concerns the relationship between alpha desynchronization and the concurrent aperiodic changes. At first glance, the coexistence of increased aperiodic exponent (suggesting increased inhibitory drive, under the E/I balance interpretation (Gao et al., 2017)) and alpha desynchronization (suggesting increased cortical excitability) appears conceptually contradictory. One resolution is to think of these as operating at different spatial scales: aperiodic changes may reflect a broader, relatively non-focal inhibitory mechanism (a large-scale “braking” signal) indicating that incoming information requires processing, while alpha desynchronization represents a more precise, spatially targeted increase in excitability over task-relevant sensory regions, potentially co-occurring with alpha synchronization over task-irrelevant areas. A broadly compatible account has been proposed by Gyurkovics et al. (2022). Interestingly, in contrast to scalp EEG studies, intracranial recordings during working memory tasks have reported a decrease in aperiodic exponent concurrent with alpha desynchronization, suggesting that both indexes increased cortical excitability at the local level (Preston et al., 2025). The divergence between intracranial and scalp findings may reflect genuine scale-dependent differences in what the aperiodic component indexes, with intracranial signals more sensitive to local synaptic dynamics and scalp EEG more sensitive to large-scale network propagation, but this remains an open and theoretically important question.

### The importance of accounting for ERPs in spectral and aperiodic estimation

A methodological consideration that deserves explicit attention is the contribution of event-related potentials to the post-stimulus frequency spectrum. Because ERP have a broadband spectral representation (Gyurkovics et al., 2022; Makeig et al., 2002; Min et al., 2007; Sauseng et al., 2007), failing to account for this activity risks attributing ERP-driven power changes to either oscillatory or aperiodic mechanisms. In the present data, ERP removal reduced the magnitude of theta-band changes, meaning that ERPs represent an additional, independent source of conflation beyond aperiodic activity. Moreover, ERP removal also reduced the observed changes in aperiodic activity, highlighting that accurate quantification of non-oscillatory dynamics likewise requires prior removal of the ERP as shown in (Gyurkovics et al., 2022). Whether the neural activity underlying the ERP is itself best characterized as additive and broadband, as a phase-reset of ongoing oscillations, or as some combination of both, remains an open and actively debated question, but one that is beyond the scope of the present paper.

### Functional significance of event-related aperiodic changes

The changes in aperiodic activity we observed overlap spatially and temporally with the effects previously attributed to theta oscillations, raising the question of whether some of the cognitive functions classically attributed to stimulus-related theta, including attentional orienting, sensory gating, cognitive control, and the updating of working memory representations (Cavanagh & Frank, 2014; Cavanagh & Shackman, 2015; Cavanagh et al., 2012; Cohen & Donner, 2013; Gyurkovics & Levita, 2021; McLoughlin et al., 2022), might instead be carried, at least in part, by changes in aperiodic activity.

This possibility is reinforced by a growing body of evidence linking aperiodic activity to a broad range of cognitive functions. Resting-state aperiodic exponent has been associated with individual differences in cognitive processing speed (Ouyang et al., 2020) and seems to dominate what is traditionally seen as “oscillatory” functional connectivity in scalp EEG (Monchy et al., 2025). Pre-stimulus aperiodic activity has been shown to predict whether a near-threshold visual stimulus will be detected (Cunningham et al., 2023), suggesting that the aperiodic component of the EEG spectrum carries meaningful information about moment-to-moment fluctuations in cognitive state. Post-stimulus aperiodic changes have been directly linked to attentional filtering performance in a cued flanker task as well, whereby larger steepening was associated with a smaller flanker interference effect (Kalamala et al., 2024). In working memory paradigms, event-related aperiodic changes have been shown to track memory load (van Engen et al., 2026), and the present data add to this picture by showing that pre-stimulus aperiodic exponent predicts individual reaction times in typically developing children, suggesting that the aperiodic background at the time of stimulus onset shapes the speed of subsequent processing.

The above findings, taken together with the fact that most stimulus-related theta-band changes appear to be accounted for by concurrent aperiodic modulation, invite a broader reconsideration of the functional architecture underlying cognitive processing. If the neural correlates of functions previously attributed to oscillatory theta are largely aperiodic in nature, then it may be that many core cognitive operations reflect the coordinated interplay of periodic and aperiodic dynamics rather than oscillatory activity alone. Under the E/I balance interpretation of the aperiodic exponent (Gao et al., 2017), this would suggest that transient, spatially specific shifts in local excitation-inhibition balance are a fundamental computational mechanism in their own right, not merely a backdrop against which oscillations operate, but an active contributor to the neural computations underlying perception, attention, and memory. Crucially, the functional significance of such local E/I shifts may depend heavily on which network is affected: the same direction of aperiodic change could have qualitatively different cognitive consequences depending on whether it occurs in prefrontal, sensory, or hippocampal circuits, but also when it happens relative to a given process, which would explain why the aperiodic exponent has been linked to such a heterogeneous range of functions across studies.

A further consideration of both methodological and functional relevance is the rotational point of the aperiodic spectrum: the frequency around which the slope pivots when the offset and exponent changes (Podvalny et al., 2015). The location of this pivot determines which frequency bands will appear to increase, and which will appear to decrease because of the same underlying aperiodic shift and therefore has direct implications for how band-limited power changes should be interpreted (Gyurkovics et al., 2022; Preston et al., 2025). In the present data, the rotational point appears to fall just above the alpha band, at approximately 13 Hz, as evidenced by the fact that theta- and alpha-band power (4-13Hz) both showed increases in the uncorrected spectrum, whereas beta and gamma power (13-40Hz) showed decreases. This pattern is precisely what would be expected from a steepening of the aperiodic slope pivoting at that frequency. It is not clear as of today if the rotational point frequency is only a mathematical artefact of how slope and offset co-vary or if it could reflect something intrinsic to the circuit, such as the characteristic time constants of dominant inhibitory interneuron populations, or the membrane integration window of the local network. If so, the pivot frequency could be informative in its own right: a shift in *where* the spectrum rotates, independent of the magnitude of the slope change, might signal a reorganization of which neural populations are governing the dynamics.

### Conclusion

Taken together, the present findings demonstrate that stimulus-related changes in aperiodic activity are a pervasive and consequential feature of the EEG signal that conventional time-frequency analyses are ill-equipped to handle. By assuming that the aperiodic background remains stable across the peri-stimulus window, conventional approaches attribute broadband spectral shifts to frequency-specific oscillatory dynamics, leading to systematic overestimation of theta and mischaracterization of beta and gamma activity. Alpha desynchronization emerges as the exception, a robust periodic phenomenon that survives and is even enhanced by aperiodic correction, providing a useful internal validation that aperiodic correction does not simply abolish all spectral effects. These findings do not imply that theta, beta, or gamma oscillations are functionally irrelevant; rather, they call for more rigorous decomposition of the EEG signal into its periodic and aperiodic constituents before drawing conclusions about oscillatory dynamics. As methods for aperiodic parameterization and the separation of periodic and aperiodic components continue to mature, their routine application in event-related paradigms will be essential for building a more accurate understanding of the neural dynamics underlying cognition and behavior.

## Competing interest

The authors declare no competing interests.

## Author contributions

T.V. and S.M. conceived the study. T.V preprocessed the data and analyzed the data under the supervision of S.M & M.Q. T.V wrote the first draft of the manuscript. S.M, M.Q, B.V and M.G edited the manuscript.

## Acknowledgment

We thank Dennis Cregin, Trinca Lecaj, and Daniella Coen for their essential contributions to participant recruitment and data acquisition. This work was supported by a grant from the Simons Foundation Autism Research Initiative (SFARI Award # 874845, SM). Support for recruitment and phenotyping of participants was provided by the Human Clinical Phenotyping Core of the NICHD funded Rose. F. Kennedy Intellectual and Developmental Disabilities Research Center (P50 HD105352, SM). The content is solely the responsibility of the authors and does not necessarily represent the official views of the National Institutes of Health.

## Ethics approval and consent to participate

This study was approved by the Institutional Review Board of the Albert Einstein College of Medicine (IRB # 2021-13433). All participants assented to the procedures and parents/guardians provided informed consent.

## Availability of data and materials

The dataset supporting the conclusions of this article is available in the ‘SFARI_EEG_multi-paradigm dataset’ repository (BIDS format), doi:10.18112/openneuro.ds006780.v1.0.0. The scripts used for preprocessing and analyses of the data are available on GitHub: https://github.com/tvanneau/Aperiodic_Theta

---

**Supplementary figure 1.**
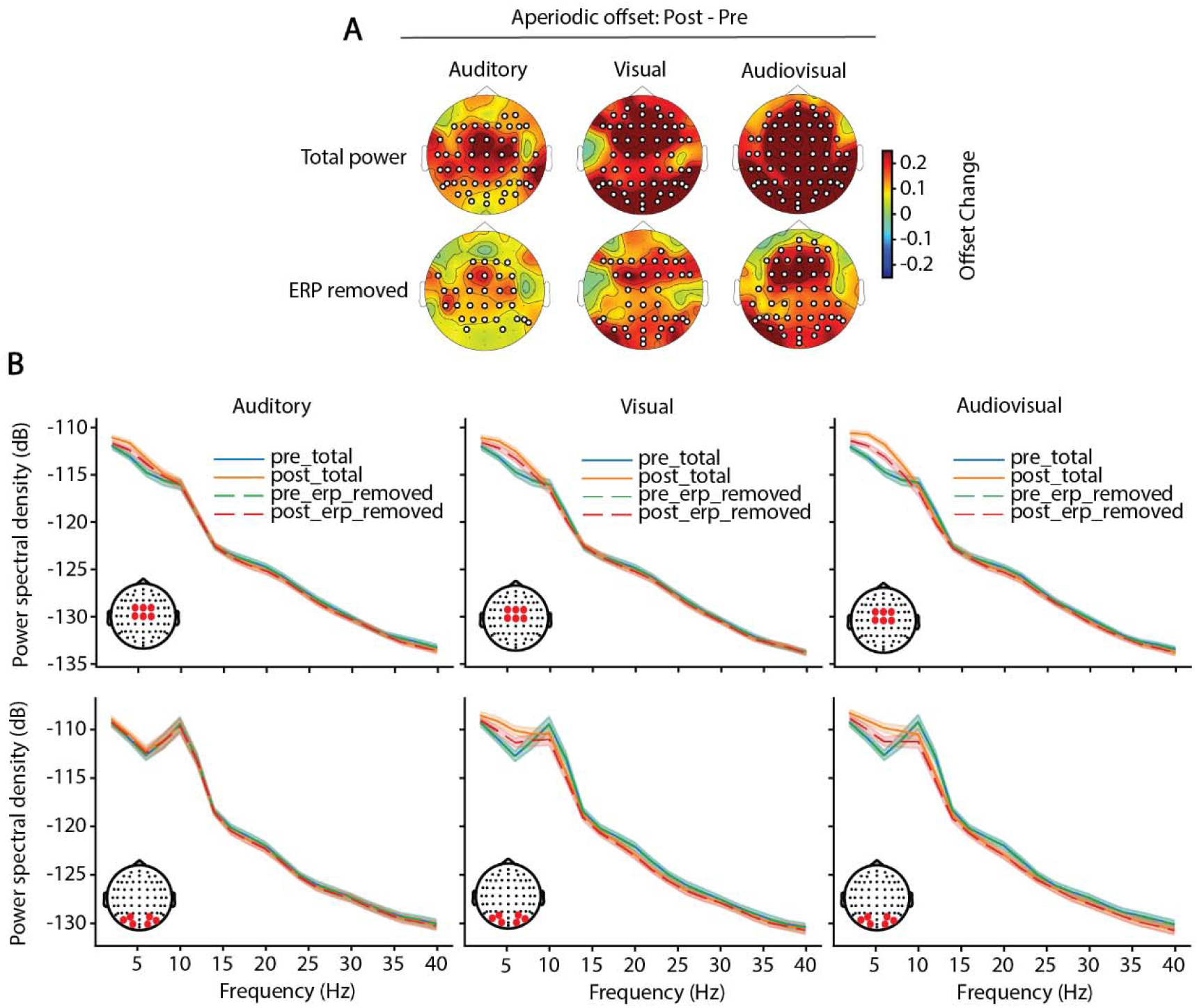
**(A)** Topographical maps of the change in aperiodic offset from pre-to post-stimulus window (post − pre) for each stimulus condition. Results are shown for the total signal (top row) and after time-domain removal of the ERP (bottom row). Black dots indicate channels belonging to significant spatial clusters identified using cluster-based permutation testing against zero change. **(C)** Power spectral density (PSD, in log space) in the pre-stimulus window (blue) and post-stimulus window (orange), shown for the total signal (solid lines) and after time-domain ERP removal (dashed lines), for auditory (left), visual (middle), and audiovisual (right) stimulation. Spectra are averaged across a fronto-central channel cluster (top row) and a parieto-occipital channel cluster (bottom row).

**Supplementary figure 2.**
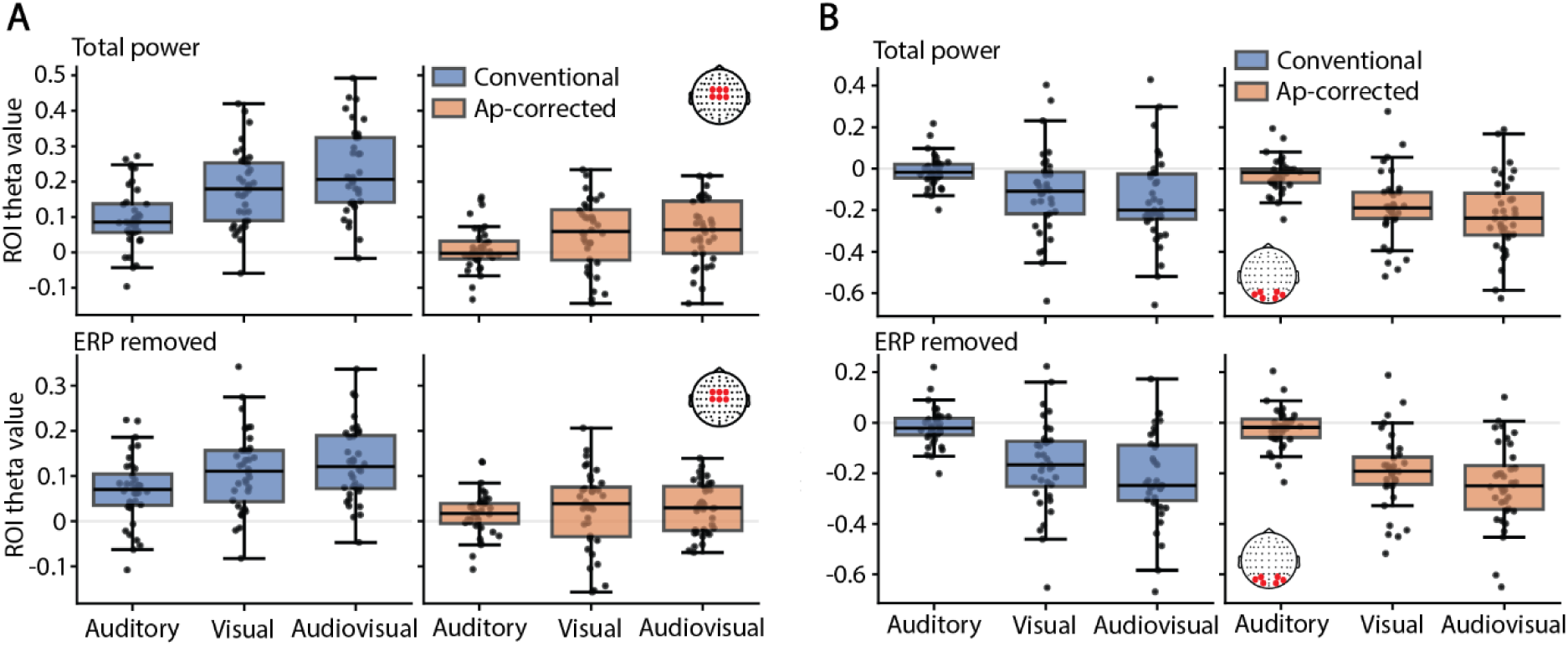
(**A**) Boxplots of averaged theta power for the conventional (left, blue) and ap-corrected method (right, orange) over a cluster of fronto-central channel for the total signal (top row) and after the removal of the ERP (bottom row). (**B**) same for alpha power.

**Supplementary figure 3.**
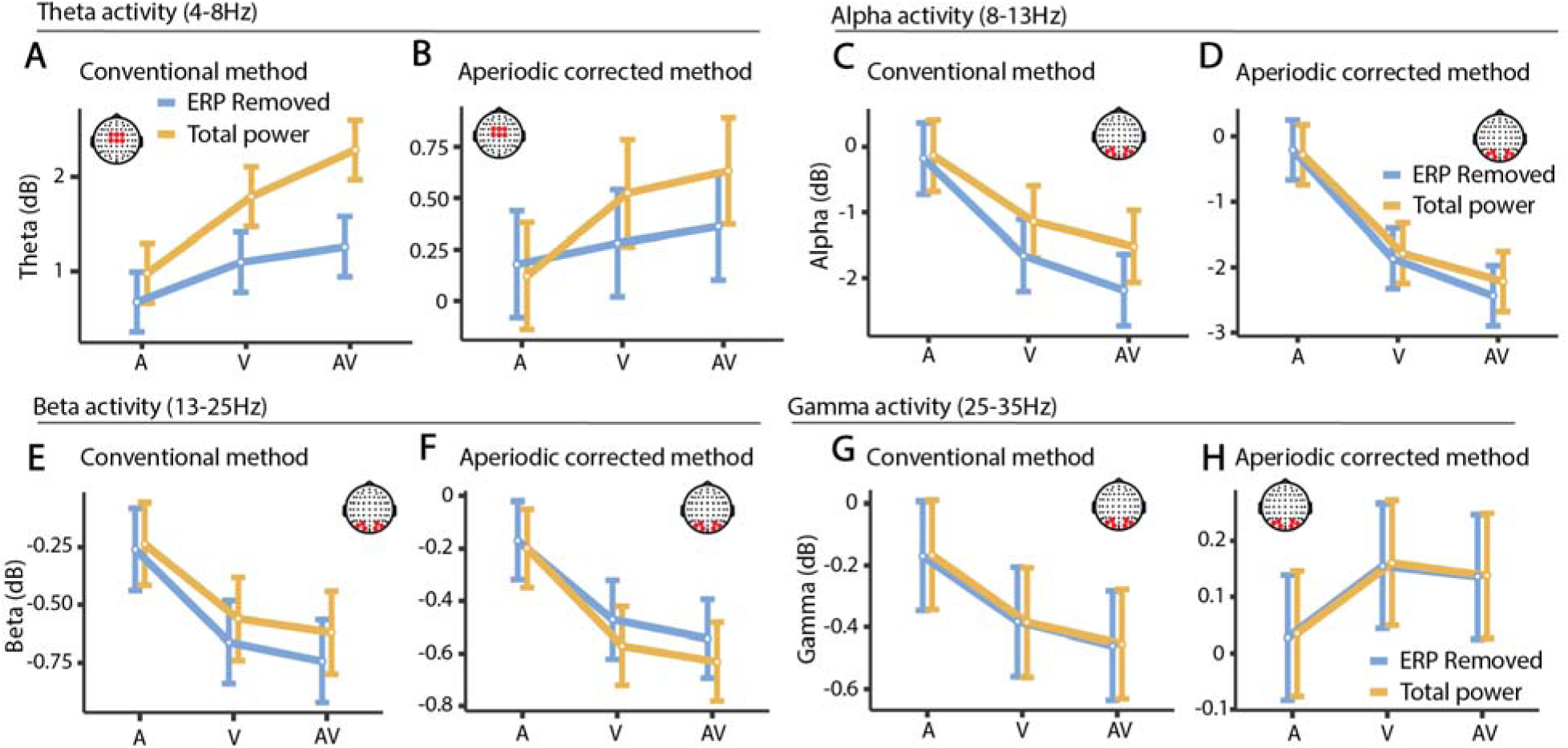
(**A**) Averaged theta power over a cluster of fronto-central channel in response to each sensory stimulation (‘A’ for auditory; ‘V’ for visual; ‘AV’ for audiovisual) for the total signal (orange) and after the removal of the ERP (blue), for the conventional method and (**B**) for the aperiodic corrected method. (**C-D**) similar then (**A-B**) but for alpha activity, (**E-F**) for beta activity and (**G-H**) for gamma activity.

**Supplementary figure 4.**
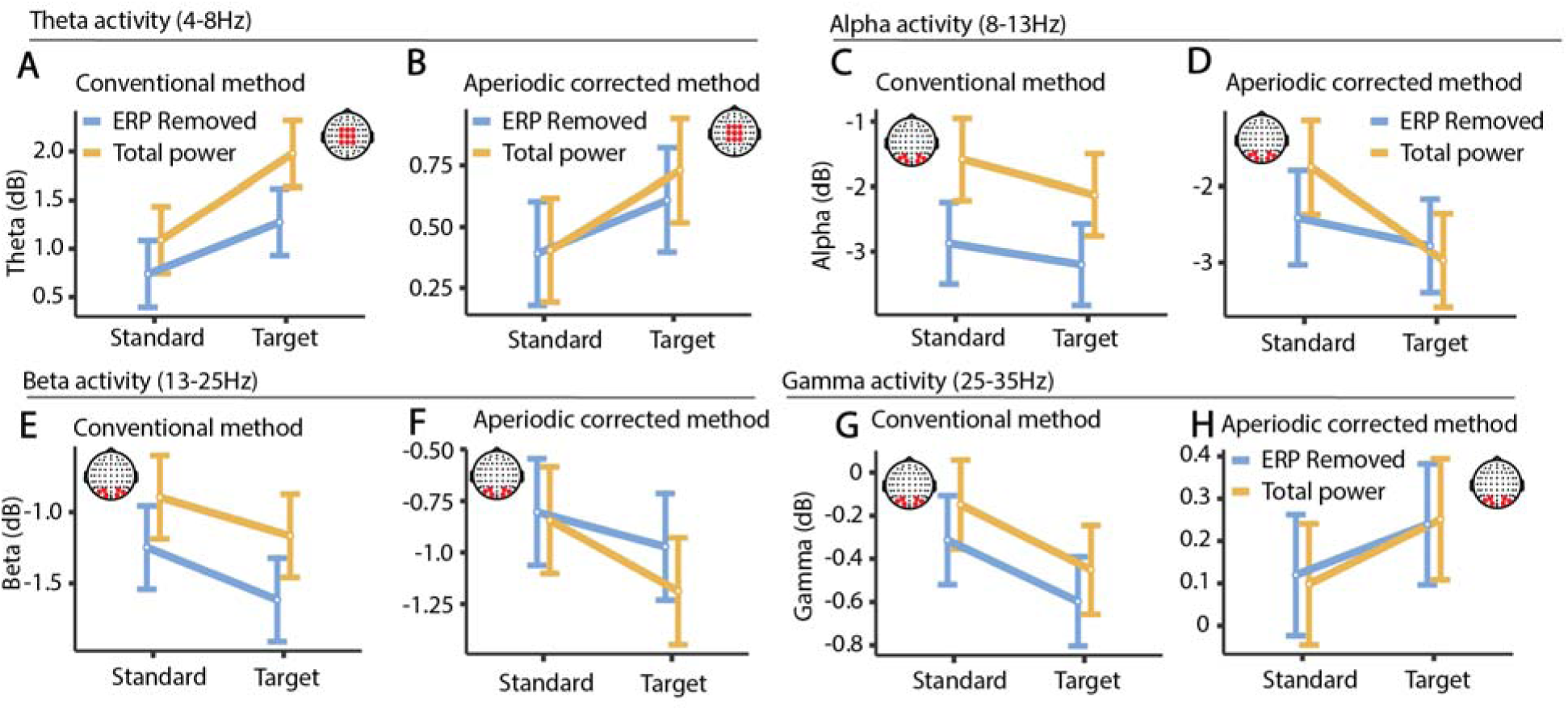
(**A**) Averaged theta power over a cluster of fronto-central channel in response to standard and target stimulation for the total signal (orange) and after the removal of the ERP (blue), for the conventional method and (**B**) for the aperiodic corrected method. (**C-D**) similar then (**A-B**) but for alpha activity, (**E-F**) for beta activity and (**G-H**) for gamma activity.

**Supplementary figure 5.**
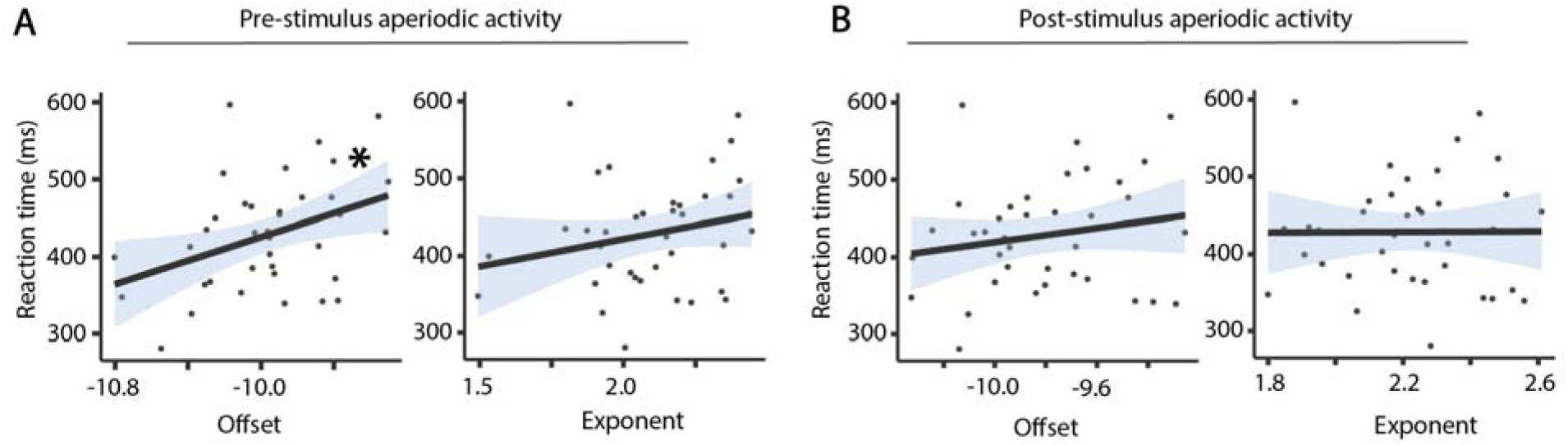
(**A**) Correlation showing the association between pre-stimulus aperiodic exponent (left) and offset (right) and reaction time during the visual oddball task, (**B**) Same analysis for post-stimulus averaged aperiodic parameters.

## Notes

### Competing Interest Statement

The authors have declared no competing interest.

